# Validating folding energy estimates as a method for variant interpretation

**DOI:** 10.1101/2025.11.09.687451

**Authors:** Clementine Elwes, Rachel Alcraft, Harris Lister, Paul Smith, David Shorthouse, Benjamin A Hall

**Affiliations:** Department of medical physics and biomedical engineering, UCL, UK; ICR, UK; Advanced Research Computing, UCL, UK; School of Pharmacy, UCL, UK

## Abstract

Interpretation of variants of uncertain significance remains a major problem in genomic analysis. Whilst statistical models can be used to predict pathogenicity, they offer no insights into the biophysical mechanism of variant action, and genomic data available for training is biased towards the subpopulations who have access. Protein misfolding has been found to act as a frequent mechanism for loss of gene or domain activity, where it is typically responsible for ∼2/3 of disease-causing variants and somatic mutations. The accuracy of energy predictions however has consistently been challenged by highly variable correlation coefficients reported from different proteins, and the unknown impact of alternative structures where available. Here we address this directly through a systematic analysis of mega-scale folding experimental results, enabled by a fully automated predictive pipeline based on FoldX. We find that whilst absolute correlation coefficients are mediocre for three highly studied proteins (0.30-0.31), the correlation coefficient alone does not capture the full predictive power of the estimates. Specifically, we find a clear linear relationship between experimental and theoretical result, with a small number of outlier residues responsible for reducing the correlation. We show that the quantitative accuracy of predictions can be improved by aggregating estimates taken from different structures, and that the problematic outlier residues can be both empirically and theoretically identified, allowing us to flag low-confidence values. Our findings not only provide a framework for identifying problematic mutations in advance but also offers new insights into potential improvements of the FoldX protocol for more accurate protein stability predictions. Our insights support the use of FoldX in computational saturation screens to support variant analysis.

## Introduction

Genome sequencing is a major tool in the study of human disease. The relentless reduction in cost over time has made these techniques increasingly widely available, having become routine in higher-income countries and starting to become available in lower and middle income countries. With increased uptake of sequencing technologies, interpretation of variants of uncertain significance is a growing problem. This is further challenged by biases intrinsic in available datasets, which have been sampled from relatively small subpopulations at this point, and the intrinsic sparsity and diversity of variants that can arise. Statistical techniques have shown great power in predicting pathogenicity but mechanistic understanding of mutation could address bias issues in existing training sets, and offers the potential for therapeutic design.

Protein misfolding is responsible for the majority of pathogenic missense mutations observed in humans (1,2), and is estimated to be responsible for a loss of activity in two-thirds of variants across multiple long-standing studies (3-5). Experimental assessments of changes in protein stability have historically been labour intensive, time-consuming, and require substantial amounts of purified protein (6). This hinders large-scale studies and the rapid assessment of protein variants. As such, computational methods are invaluable tools in high-throughput studies involving numerous protein variants.

A wide range of tools are now available for calculating folding energies of protein structures, with FoldX (7) and Rosetta (8) leading the field. FoldX specifically is one of the most widely used tools for predicting the effects of mutations on protein folding and stability (9), and has been found to offer similar predictive power to Rosetta at substantially lower computational cost (10). FoldX stability calculations have been used to assess the impact of various disease-linked mutations on the stability of the protein rhodopsin in the retina (11), and missense mutations in E-cadherin on the protein’s stability and its role in the hereditary diffuse gastric cancer (12). FoldX has also been effectively used to study the impact of mutations in viral proteins, particularly in the context of SARS-CoV-2. One study combined molecular dynamics simulations with FoldX to predict escape mutations in the virus’s receptor-binding domain, finding a positive correlation between predicted binding affinity changes and experimentally observed escape fractions (13). Another study assessed the effects of mutations on the binding interactions between the SARS-CoV-2 spike protein and neutralising antibodies, aiding in the identification of mutations that could impact vaccine efficacy (14).

Whilst FoldX has been successfully applied to a wide range of different biological problems, there remains ongoing questions regarding its predictive power and therefore its suitability for assessing proteins generally. This is highlighted by the wide range of correlation coefficients reported in the literature, for example from 0.2 for the engineered Flavin mononucleotide based Fluorescent Protein (15) to 0.8 for a large set of mutations across multiple proteins (7). Related to this is how the choice of initial structure influences calculations. This has been studied for different structures of COVID spike protein, revealing variation between estimates taken from different structures and the need to aggregate data to establish a consensus (16). Resolving and validating FoldX predictive performance at scale across multiple proteins is fundamental to its adoption in variant interpretation.

Here we present a systematic analysis of FoldX predictions through computational saturation screens applied to a mega-scale study of protein folding stability from Tsuboyama et al (17). We focus primarily on the proteins for which Tsuboyama et al demonstrated high agreement (Pearson r >0.75) between their energy estimates and previously published data. This dataset of over 1000 well-validated substitutions in seven proteins with large numbers of experimentally determined structures allows us unprecedented validation of predictions generated by the tool. We find that the presence of outliers substitutions, which make up a small proportion of the overall estimates, distorts the correlations between predictions and measurements, obscuring a clear linear trend. When comparing across alternative structures for a single protein we find estimates for single substitutions are generally consistent, with unimodal distributions well described by the median. Using elastic network modelling we find that outlier substitutions are associated with residue positions that are tightly constrained within structures, leading to overestimates. Finally, across the full set of c. 200 proteins for which Tsuboyama et al derived experimental folding energies, we find taking the median FoldX energy prediction across available structures for a protein yields correlations with experimental energies approaching the upper limit set by experimental reproducibility (r ≈ 0.75).

## Materials and Methods

### Mutein pipeline for automated FoldX ΔΔG calculation

Following the method originated in Shorthouse et al (10) and adapted to a high-performance computing cluster for Hall et al (18), FoldX was used to calculate the ∆∆G values of mutations on a Sun Grid Engine computing cluster. For each gene of interest, the pipeline retrieved all PDB structures from the primary accession code using the UniProt API (19), and additionally the AlphaFold PDB structure (20). For every PDB structure, a parallel FoldX RepairPDB command was run 5 times for steric clash minimisation and optimisation of residue orientations. With further parallelisation, FoldX PositionScan was run on every residue in the relevant chains, for every possible amino acid mutation. The default FoldX settings were used for RepairPDB and PositionScan. The resulting protein residues are aligned to the gene positions resulting in ∆∆G for every possible mutation on all relevant PDB structures for the gene. Code for performing simulations is available at Zenodo (21). All data is included in supplementary information.

### Gene alignment

Structures downloaded by the Mutein pipeline frequently are not aligned to the canonical amino acid sequence for the gene as defined by UniProt. Differences can include first residues (for example, due to the construct used for structural analysis), residue numbering, excised loops, or natural variants. To ensure the structures are aligned to the reference protein sequence, we extract from the datasets the sequence used in the FoldX pipeline for each protein structure and the sequence used in the experimental dataset.

Our alignment approach is multi-step. Structures with dataset sequences exactly matching the reference gene sequence in Uniprot were identified. For structures with dataset sequences exactly matches the PDB sequence, residue level mappings to the reference gene sequence were obtained using the PDBe API. For dataset sequences that did not match, a smart k-mer search was used to identify exact subsequences within the reference gene sequence, which served as anchors for finding the alignment. For homomeric proteins, alignments from a representative chain were propagated to all identical chains.

We further take account of amino acid substitutions in the experimental structure compared to the reference when returning the ΔΔG of a substitution elsewhere. Assuming a thermodynamic cycle, we add the ΔΔG of mutating back to the reference sequence to the substitution being studied for that structure so that all energies under study are relative to the same reference sequence. All code used in this study is available as SI.

## Results

### Analysis of PIN1 reveals underlying relationship between theoretical and measured results

PIN1 is a nuclear localised protein responsible for the control of multiple cellular processes, including cell cycle progression and the DNA damage response. It acts as an enzyme catalysing conformational transitions in proteins, initiating changes in protein structure, stability and function (22). It consists of two domains connected by a disordered region; a WW domain that mediates specific protein-protein interactions, and a PpiC domain responsible for catalysis. Of proteins studied in the high-agreement experimental dataset, PIN1 has the most available structures (Table 1), and so was selected for initial analysis. All structures were found to have different residue numbers (Fig 1A), so were realigned to the UniProt reference sequence before being correlated with experimental data (Fig 1B, S1, materials and methods).

**TABLE I.** Problematic Residues and Mutations.

| Protein | (1)<br>Total<br>Residues | (2)<br>Total<br>Mutations | (3)<br>Problem<br>Residues | (4)<br>Problem<br>Mutations | (5)<br>Problem<br>% Outliers |
| --- | --- | --- | --- | --- | --- |
| PIN1 | 32 | 60 | 4 | 4 | 71% |
| Spg | 54 | 886 | 4 | 19 | 70% |
| FYN | 14 | 80 | 4 | 5 | 79% |
| RplI | 30 | 42 | 2 | 4 | 77% |

**Figure 1.**
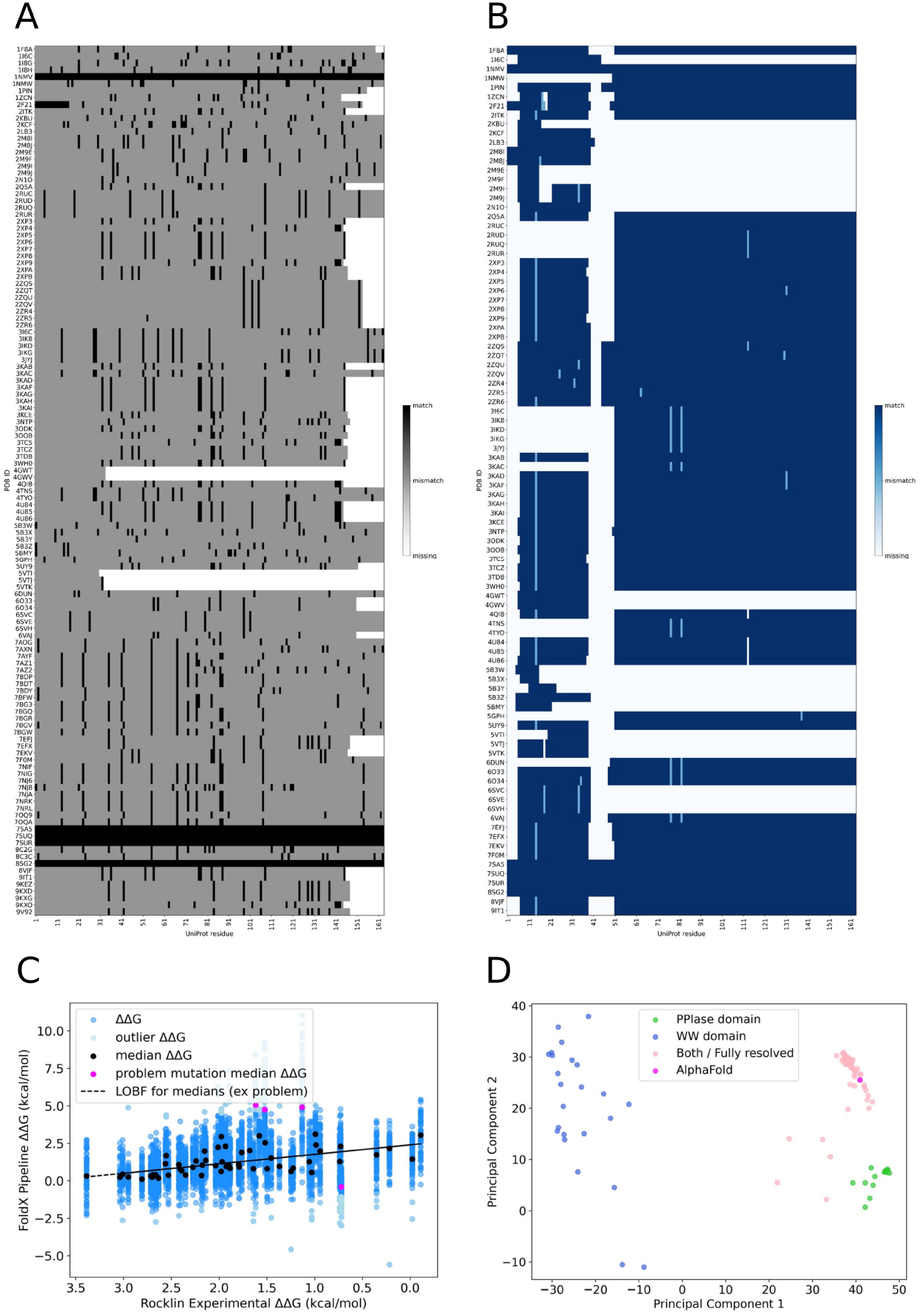
Theoretical estimates of PIN1 variant misfolding reveal underlying predictive relationship. A) Before alignment heatmap PIN1 residue matches between PDB structures and the reference sequence for PIN1. Black is a residue match, grey is a mismatch, white is no residue. B) After alignment heatmap of residue matches between FoldX pipeline structures and the reference gene sequence for PIN1. Dark blue is a residue match, pale blue is a substitution, and blank is either a mismatch or no residue. For PIN1 we can see that residue positions 40-50 are a loop that isn’t historically resolved easily, other than in 4 relatively new structures. C) Scatter plot of predicted and measured change in energy of folding (ΔΔG) with linear regression analysis, outlier identification and median energies. When problematic residues are discounted, median energies tighten the linear trend. D) PCA projection of top two components with domain colouring for PIN1, with each structure normalised independently so that PCA reflects variation within local domains, revealing the relationship between experimental construct and folding energy prediction. PIN1 has two resolved domains, the WW domain which is residues 1-38 and the PPlase domain which is residues 50 up to 163. Most structures do not resolve the loop region in between. AlphaFold energy estimates cluster with structures of the full structure.

An initial analysis of theoretical estimates and measurements suggests a weak relationship between the two, with a Pearson correlation coefficient equal to 0.30. Visualisation of the scatter plot however reveals a clear linear trend, punctuated by a small number of poorly predicted substitutions (Fig 1C). A linear regression applied to the data confirms the observation, and allows us to highlight outlier substitutions as those that are more than two standard deviations away from the line of best fit. This shows that four substitutions from four residues make up 71% of all the outliers.

The outsized role of individual substitutions raises questions on the nature and origin of variation between different estimates taken from structures that are ostensibly the same. To explore this question for individual substitutions we tested the shapes of the distributions. We found using Hartigan’s dip test that the distribution of estimates was unimodal for almost all substitutions (with only substitution (T29D) less than a multiple test corrected p-value threshold of 0.05). Application of the D’Agostino-Pearson test revealed that a fifth of these unimodal distributions were not normally distributed (12/60 non-normal), and had a skew apparent from visualisation of individual distributions. The median of these distributions was found to adequately represent the peak of this distribution, and when the median is plotted against experimental energies this is observed to tightly follow the trend of the linear regression resulting in an improved Pearson correlation coefficient of 0.61. These observations raise the question of how much do measures of ΔΔG vary between different structures for well behaved substitutions. For PIN1, a bootstrapping analysis of non-outlier substitutions reveal 95% confidence intervals of -0.27 and +0.35 kcal/mol. Outlier substitutions have a notably broader variance than other substitutions, suggesting that problematic residues can be identified from this distribution and flagged as low confidence predictions.

Finally, to explore the nature of variance across the complete dataset, we performed a principal component analysis of all measured energies. A biplot shows a separation of structures based on the specific experimental construct used in determining the structure (Fig 1D). This has the further effect of separating NMR structures from crystal structures, as a result of using smaller constructs. Given that AlphaFold models are folded using forcefields trained from potentially all available structures, we examined the energies from this model finding that it clustered with full length constructs, but the energies had a weaker correlation with experiment in isolation than the median energies.

### Expansion to a larger protein cohort confirms underlying patterns in prediction

The observations in PIN1 show that FoldX was capable of producing accurate predictions of folding energies, and suggested a process for handling the energy estimates taken from a large number of structures. To validate these insights, we first analysed the predicted energies for FYN and Spg, the proteins from the mega-scale dataset with the next two largest numbers of structures. *FYN* encodes a kinase in the src family (23), whilst *spg* in Streptococcus sp. group G encodes immunoglobulin G-binding protein G (24).

Table 1. Distribution of problematic substitutions across proteins under study. “Problem” residues or mutations are defined by those with estimates more than 2 standard deviations away from the line of best fit. A small minority of residues contribute disproportionately to outlier observations, defined as being more than two standard deviations from the line of best fit. THO1, CPA2, PRPF40A are excluded from the analysis as there are insufficient datapoints to identify outliers.

For some SPG and FYN structures, individual substitutions relative to the reference sequence have extremely large ΔΔG values. When applying the thermodynamic-cycle correction, these values generate substantial global offsets, resulting in strongly negative rebased ΔΔGs for all positions in that structure. We retain these structures in the linear regression and correlation analyses to preserve a common thermodynamic reference. However, the identification of problematic mutations in SPG and FYN is based on the large positive ΔΔG outliers observed in Fig. 2, rather than on the absolute magnitude of the rebased energies.

**Figure 2.**
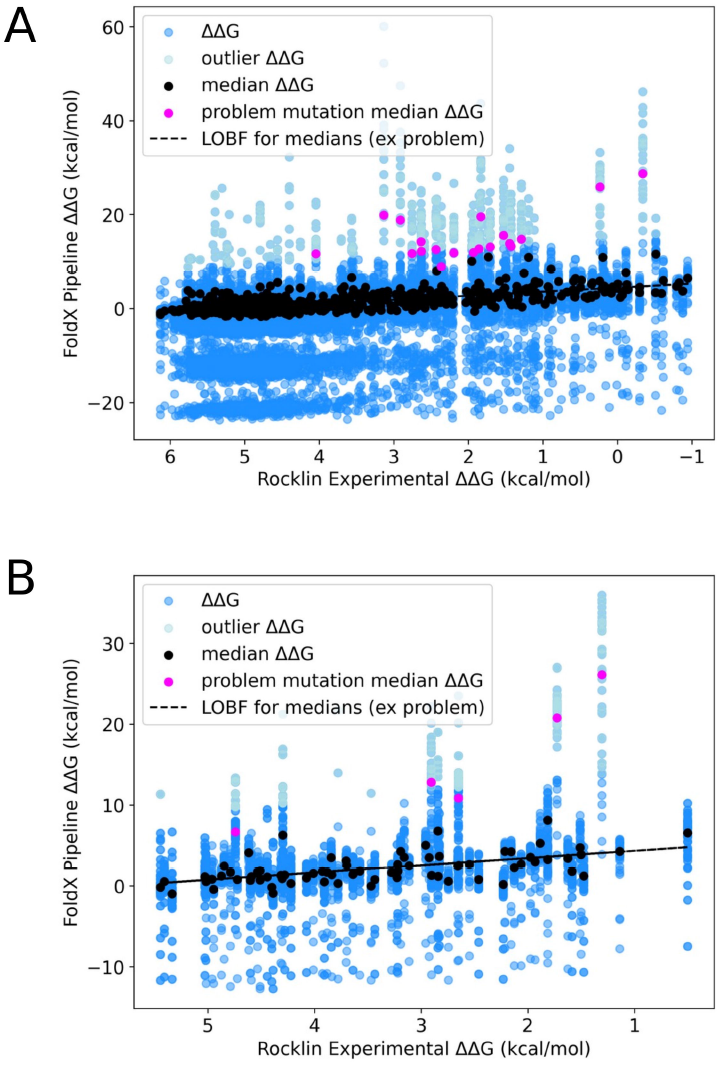
Linear relationship between theoretical estimates and measured energies is observed for (A) SPG and (B) FYN. Consistent with observations from PIN1, predictions and measurements have a linear relationship, with a handful of outliers reducing the correlation. Visualising the median estimates for each substitution we see that they cluster tightly to the linear trend of the whole cohort. Each gene is found to have a slightly different slope. Points falling far below the line of best fit correspond to structures with extreme negative rebasing caused by large offsets in the thermodynamic-cycle correction.

Nevertheless, the same patterns observed in PIN1 were recapitulated (Fig 2) for FYN and Spg. Both sets of predicted energies follow a line of best fit, with a stronger correlation found between the medians (0.72 for Spg, 0.58 for FYN) than the complete set of predicted energies. In each case a small number of substitutions contributed disproportionately to the positive outlier energies observed (Table 1). When considering only non-outlier residues, we find comparable 95% confidence intervals across different structures to those see in PIN1 (Spg -0.597, +0.479 kcal/mol, FYN -0.528, +0.478 kcal/mol). These trends were further confirmed when expanding beyond the three initial proteins to the complete datasets (Fig S2).

While the automated tool facilitates alignment of all structures of all proteins in the experimental dataset, the level of analysis going forward is limited by the representation of each protein in the high-agreement experimental dataset. For three of the ten proteins (SUCB, YAP1, VIL1) there are only two or three single variants in the experimental assay and hence analysis of these proteins is not taken further.

Table 2. Slope and standard error of the slope of the line of best fit calculated from measured and predicted misfolding energies.

**TABLE II.** Linear Slope Coefficients.

| TABLE II<br>LINEAR SLOPE COEFFICIENTS |  |  |  |  |
| --- | --- | --- | --- | --- |
| Protein | (1)<br>Structures | (2)<br>Mutations | (3)<br>All estimated $\Delta\Delta G$ | (4)<br>Median estimated $\Delta\Delta G$ |
| **PIN1** | 79 | 60 | 0.60 | 0.63 |
| *St. error* |  |  | 0.03 | 0.11 |
| **Spg** | 28 | 886 | 1.5 | 0.86 |
| *St. error* |  |  | 0.023 | 0.028 |
| **FYN** | 38 | 80 | 1.2 | 0.90 |
| *St. error* |  |  | 0.059 | 0.15 |
| **CPA2** | 2 | 20 | 0.93 | 0.97 |
| *St. error* |  |  | 0.25 | 0.23 |
| **PRPF40A** | 4 | 47 | 0.46 | 0.33 |
| *St. error* |  |  | 0.16 | 0.19 |
| **RplI** | 5 | 42 | 0.14 | 0.55 |
| *St. error* |  |  | 0.14 | 0.11 |
| **THO1** | 3 | 28 | 0.55 | 0.64 |
| *St. error* |  |  | 0.18 | 0.14 |

The common trends observed for each protein in isolation raises the question whether the trends hold when all predictions aggregated together (Table 2). Examination of the trend lines for different datasets shows that each line of best fit appears to have a distinct slope and intercept. It should be noted however that 95% confidence intervals are considered, all slope coefficients form a continuous chain of overlaps across the proteins and hence there is no suggestion they follow statistically different models.

After processing the FoldX pipeline ΔΔG estimates by removing problem substitutions where there are sufficient datapoints to identify them, and aggregating across structures using the median, correlation coefficients are calculated for each protein (Table 3). The correlation between the two experimental datasets in column 3 represents an upper bound on the agreement a theoretical prediction can achieve, as it reflects the intrinsic noise variability between independent experimental assays. The correlations of the FoldX pipeline with each experimental dataset approach the upper bound quite closely for many proteins while for other proteins with fewer datapoints correlations can be lower (PRPF40A in particular). Overall, the results demonstrate that the FoldX pipeline captures a substantial portion of the reproducible signal in experimental data across multiple proteins and justifies using FoldX to predict misfolding energies.

**TABLE III.** Pearson Correlation Coefficients.

| Protein | (1) | (2) | (3) |
| --- | --- | --- | --- |
|  | PipelineTheor-RocklinExp | PipelineTheor-PrevExp | RocklinExp-PrevExp |
| PIN 1 | 0.61 | 0.55 | 0.84 |
| spg | 0.72 | 0.68 | 0.81 |
| FYN | 0.58 | 0.45 | 0.75 |
| CPA2 | 0.70 | 0.49 | 0.76 |
| PRPF40A | 0.24 | 0.37 | 0.85 |
| rplI | 0.66 | 0.66 | 0.90 |
| THO1 | 0.68 | 0.70 | 0.95 |

Table 3. Pearson correlation coefficients between the FoldX pipeline theoretical predictions and experimental measurements. Correlations for PIN1, spg, FYN and THO1 are calculated after removing problem substitutions identified in Table 1 and aggregating predicted ΔΔG across structures by the median. For CPA2, rplI and PRPF40A, as there are insufficient datapoints to identify outliers, correlations are based on median aggregated energies. Column 3 reports correlations between two independent experimental datasets, representing an approximate upper bound for correlations achievable by theoretical predictions.

### Outlier predictions arise from highly constrained residues

Whilst analysis of the whole cohort reveals fundamental trends that enable predictions, outlier substitutions are a clear problem for a small number of residues. Understanding the origin of these problematic substitutions may suggest methods to address the issues, or flag hard to predict residues at the point of calculation to users. We therefore studied the origins of the outlier substitutions.

We first sought to conclude whether all substitutions of single residues are poor predictors, or if specific substitutions are problematic. For all proteins other than Spg we cannot draw conclusions on this as the high-agreement Tsuboyama dataset does not have experimentally observed ΔΔG for all substitutions at a particular residue position (Table 1). For PIN1, FYN and RplI it appears that if a residue is problematic then all substitutions at that point are poor. However, given only select substitutions at these residues are represented in the Tsuboyama dataset this is arguably a sample effect. For Spg however, all substitutions are evenly distributed in variants (Fig S3) and so we can look at the distributions of substitutions in the outliers. Substitutions in the outliers are dominated by bulky aromatic amino acids tyrosine (Y) and phenylalanine (F) as well as bulky polarised amino acid histidine (H) with the count of tyrosine and phenylalanine 4-times and the count of histidine 2-times that of the other amino acids of substitution (Fig S3). Structures that exhibit extremely negative rebased ΔΔGs due to large offsets in the thermodynamic-cycle correction were excluded from analysis with variance in ΔΔG. With problem mutations categorised by high variance in ΔΔG values we can also identify problem residues across the protein sequence without experimental measurements, increasing the size of our dataset and enabling deeper analysis. As Fig 3A illustrates for PIN1, Spg and FYN the feature of a few residues with large variances in ΔΔG values observed when validating the FoldX pipeline with the experimental dataset above (Fig S4A,C) is a feature for the set of protein structures. Most amino acid substitutions (except alanine (A) and glycine (G)) contribute to the spikes in energies in Fig 3A, with bulky aromatic and basic residues dominating the distribution (Fig 3B).

**Figure 3.**
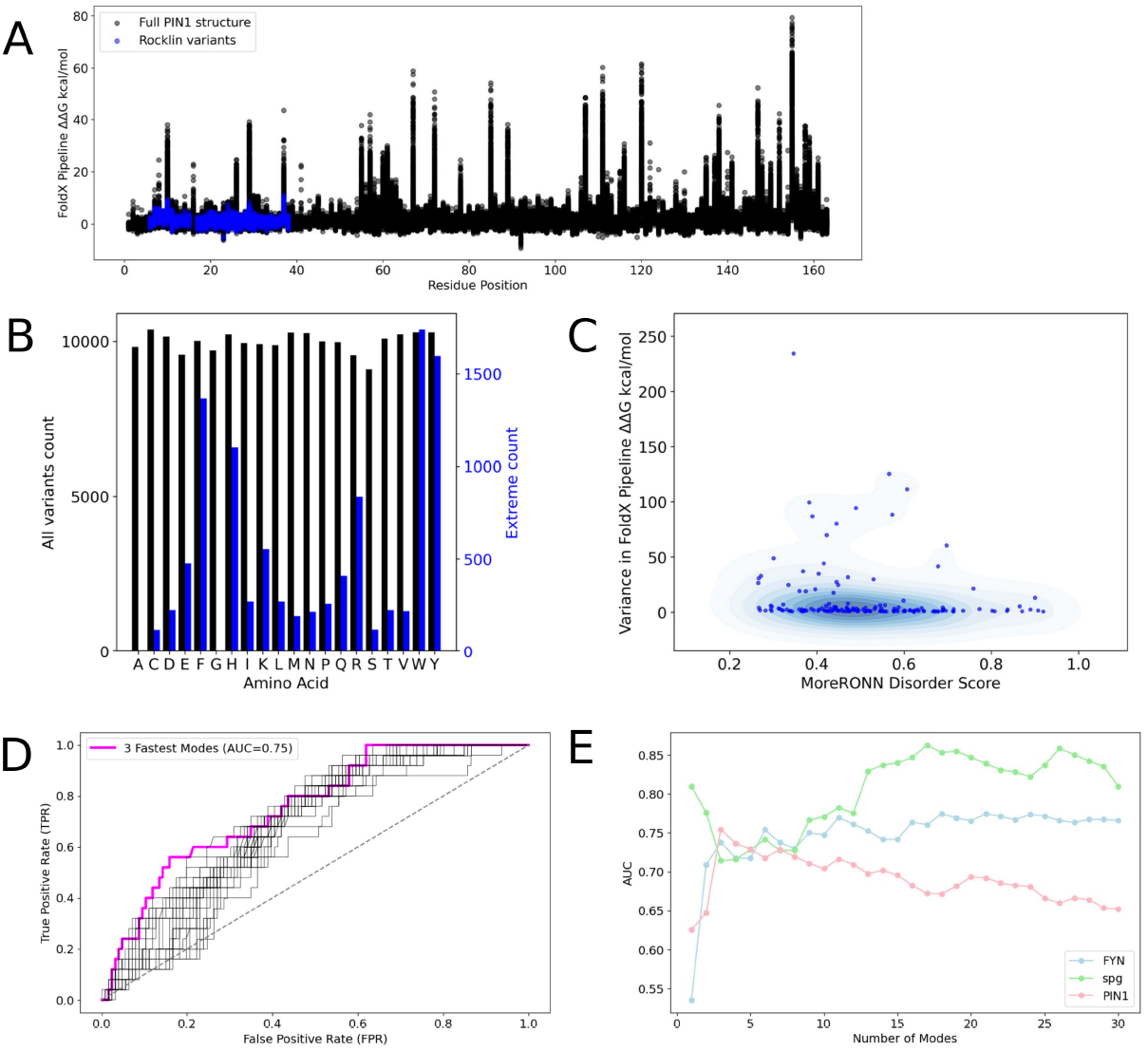
Outlier estimates arise from tightly packed residues. A) Visualisation of energy estimates from all predictions reveals that all substitutions to single residues (rather than single substitutions) share outlier estimates. B) Energy spikes in A (>10 kcal/mol) are associated with most amino acid substitutions (except alanine and glycine) and are dominated by bulky aromatic and basic residues. C) Outlier estimates are not found in disordered areas as predicted by MoreRONN. Scatter plot of energy variance against score shows no clear relationship. D) Gaussian network modelling of tightly restrained residues is predictive of outliers in PIN1. ROC curve plots false and true positive rates for different summed squared fluctuations of residues in top 3 modes (AUC=0.75) E) The number of modes needed to predict outlier residues accurately varies between genes. The multiple black curves represent the ROC curves for the first fastest mode up to the 30 fastest modes while the mode in magenta is the mode that maximises the area under the curve (AUC).

At this point we hypothesise whether from a full protein perspective you see higher energies in regions of more disorder as for example for PIN1 the most structured well-defined part of the protein appears to be the WW domain (residue positions 1-38). This would be expected to arise if flexible loop regions are poorly packed in the minimised structure. To test this we use MoreRONN scores as a predictive score of disorder. MoreRONN gives a value for every residue in the protein so we can plot the variance in ΔΔG values against the disorder score (Fig 3C). It appears for PIN1 and FYN that more ordered areas have the highest variance in energies, while the reverse is the case for Spg where higher variances are found in the more disordered regions. While permutation tests suggest they are statistically significant individually for PIN1 and Spg these relationships are small in real terms with correlation coefficients less than 0.2. In addition, a Kruskal-Wallis H-test to test if they are from different distributions is not statistically significant. We also note that for Spg only small regions of the protein are resolved and these overlay solely with regions of disorder (Fig S4A), whereas the complete protein for PIN1 and FYN is resolved across structures. As such it is also possible the relationship observed in Fig 3C – suggesting higher variances in ΔΔG values are found in more disordered regions of Spg - is largely a sampling artifact.

Based on these observations, we hypothesised next that problematic residues that are harder for FoldX to predict ΔΔG are in tightly structured regions of the protein. To investigate this we use an Elastic Network Model to find the mode of motions for the protein from the static structure, examining the fastest modes of motion (25,26). These are small localised high frequency vibrations in incredibly tight energy wells, and are believed to represent highly constrained regions. We initially examined the 10 fastest modes for PIN1 and found a significant result using a Chi-squared test (p value = 0.017), suggesting residues with a high variance in energies are not independent of residues in the 10 fastest modes. We next looked to determine how many fast modes of motion best predict residues with high variance in energies. We took from the 30 fastest modes for each protein and plotted the receiver operating characteristic (ROC) curve, to allow us to identify the optimal number of fast modes to consider for each protein. Fig 3D and S4B,D show the ROC curves for PIN1, Spg and FYN. The area under the curve of the optimal number of fastest modes is 0.75 for the 3 fastest modes for PIN1, 0.86 for the 17 fastest modes for Spg and 0.77 for the 18 fastest modes for FYN. There is however a notable lack of consensus across the proteins for the best number of modes in the classifier (Fig 3E). Together these show that the problematic predictions arise from substitutions to keystone residues in maintaining the protein fold that are not adequately refolded or minimised after computational mutation.

### FoldX is a good predictor when expanded to full extended gene set

To validate the observations from the initial set of proteins, we applied the same pipeline to the expanded set of ∼ 200 proteins for which Tsuboyama et al derived experimental folding energies. Of these, 58 proteins have sufficient assay coverage and experimentally determined structures to be included in the analysis. Within individual proteins, the FoldX pipeline recapitulates relative mutational effects and captures a substantial fraction of the reproducible signal in the experimental data. As shown in Fig. 4, aggregating ΔΔG estimates by the median across structures for each protein improves linear correlation coefficients toward the upper bound set by experimental reproducibility (r ≈ 0.75).

**Figure 4.**
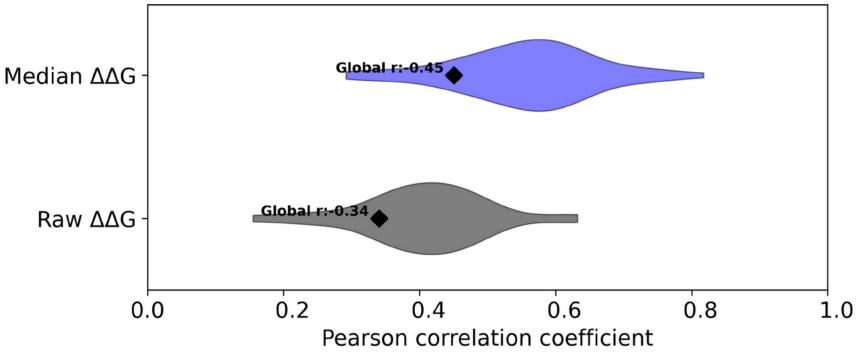
FoldX captures linear trends in ΔΔG within individual proteins. Aggregating theoretical ΔΔG estimates by the median for protein structures in the expanded Rocklin dataset with at least 6 residues in the assay and 5 experimentally derived structures (58 structures of the 206 in the dataset), improves the linear correlation coefficients for each protein towards the upper limit set by experimental reproducibility (r ≈ 0.75). Pooled correlation across the 58 proteins is poorer - the global Spearman correlation for median ΔΔG energies is 0.45.

If different proteins exhibit distinct ΔΔG scales and slope coefficients, pooled linear correlations would be strongly attenuated. We therefore assess global agreement using Spearman rank correlation. This yields a reduced correlation of r=0.45. While FoldX performs well when assessing which substitutions are more destabilising within a single protein it does not reliably reproduce absolute ΔΔG magnitudes across proteins.

## Conclusions

Protein misfolding is a common mechanism underpinning gene dysfunction through mutation. Accurate and rapid prediction of the impact on protein stability of an amino acid substitution associated with disease allows us to predict pathogenicity and potentially propose new therapeutic interventions. It also offers the opportunity to separate distinct mechanisms of gene dysfunction in heterogeneous datasets such as tumour sequencing (27). Our work presented here is a major validation of FoldX as a tool for prospective calculation of misfolding energies. It rationalises the apparent wide variation in measured correlation coefficients across different proteins, and establishes that the approach is sound for both qualitative and quantitative prediction. We have shown that small numbers of positions are poorly predicted, and that these are in themselves predictable, allowing us to propose strategies to manage them. Our work allows us to estimate the confidence intervals due to variations between structures, finding +/-0.27 to 0.60 kcal/mol, allowing us to estimate confidence for proteins with fewer or single structures available. Our findings that outlier residues distort analyses further resolves contradictions between studies that reported poor correlation between experimental energies and predictions, alongside studies that showed strong predictive power for individual proteins. These outliers can severely reduce correlations whilst only affecting a small proportion of substitutions. The work also reveals previously unreported insights from inter-protein and inter-structure comparisons. These tantalisingly suggest that different relationships between predicted and measured energies may be associated with different proteins, and that variation between structures of the same protein are principally dependent on construct. Together this underlines the influence of protein environment (including experimental conditions) in determining the fold of a structure.

Despite the new insights gathered, we acknowledge there are limitations in the data used for validation and the implications for our study. The initial dataset comprised small well-folded proteins, with no oligomers or cofactors and did not represent common protein types including multimeric proteins, DNA binding proteins, and membrane proteins. We expanded the analysis with a dataset that includes transcription factors, chromatin-associated proteins, signalling kinases and multimeric enzyme complexes. However strongly cofactor-dependent systems or membrane-embedded proteins, that would be expected to potentially present unique problems that may challenge the forcefield or require alternative options to be selected in the running of FoldX, remain underrepresented. Nonetheless, our study represents a comprehensive validation of this approach and future experimental datasets will extend our understanding of prediction of misfolding.

Our work also highlights practical issues that may undermine prediction. Estimates are worse where there are few structures available. Outliers tend to be overestimates, and arise from poor repacking in the process of minimising the structure. Future approaches may benefit from using protein dynamics to address both issues. Low-computational cost simulation methods such as CONCOORD or BioEmu could be used rapidly generate large numbers of alternative structures (28,29). Similarly, whilst AlphaFold models did not generate better predictions, refolding each model after mutation with AlphaFold or BioEmu may minimise the impact of poor repacking post mutation. The fundamental trade off here is that ensemble based approaches like Rosetta increase computational time required, and where it has been studied has not greatly altered predictions, and addressing this will require further work.

The results presented here justify the expansion of FoldX adoption and other computational saturation screen based approaches to prediction of changes to protein function through mutation (30,31). In addition to the benefits of mechanistic understanding, rapid and accurate estimation of biophysical properties could offer a new set of features to decorate sequencing datasets and support machine learning approaches. In this light, our study further justifies the future collection of pre-calculated databases of energies of misfolding as a complement to other public scores of pathogenicity.

## Acknowledgments

Thanks to Matthieu Bougueon for comments on the manuscript. BH gratefully acknowledges support from CRUK (CDEPIL-Jan24/100032).

## Data and code availability

The mutein pipeline is made available through Zenodo (21). All other code and data are shared as supplementary materials.

## Figure Legends

**Supplementary Figure 1.**
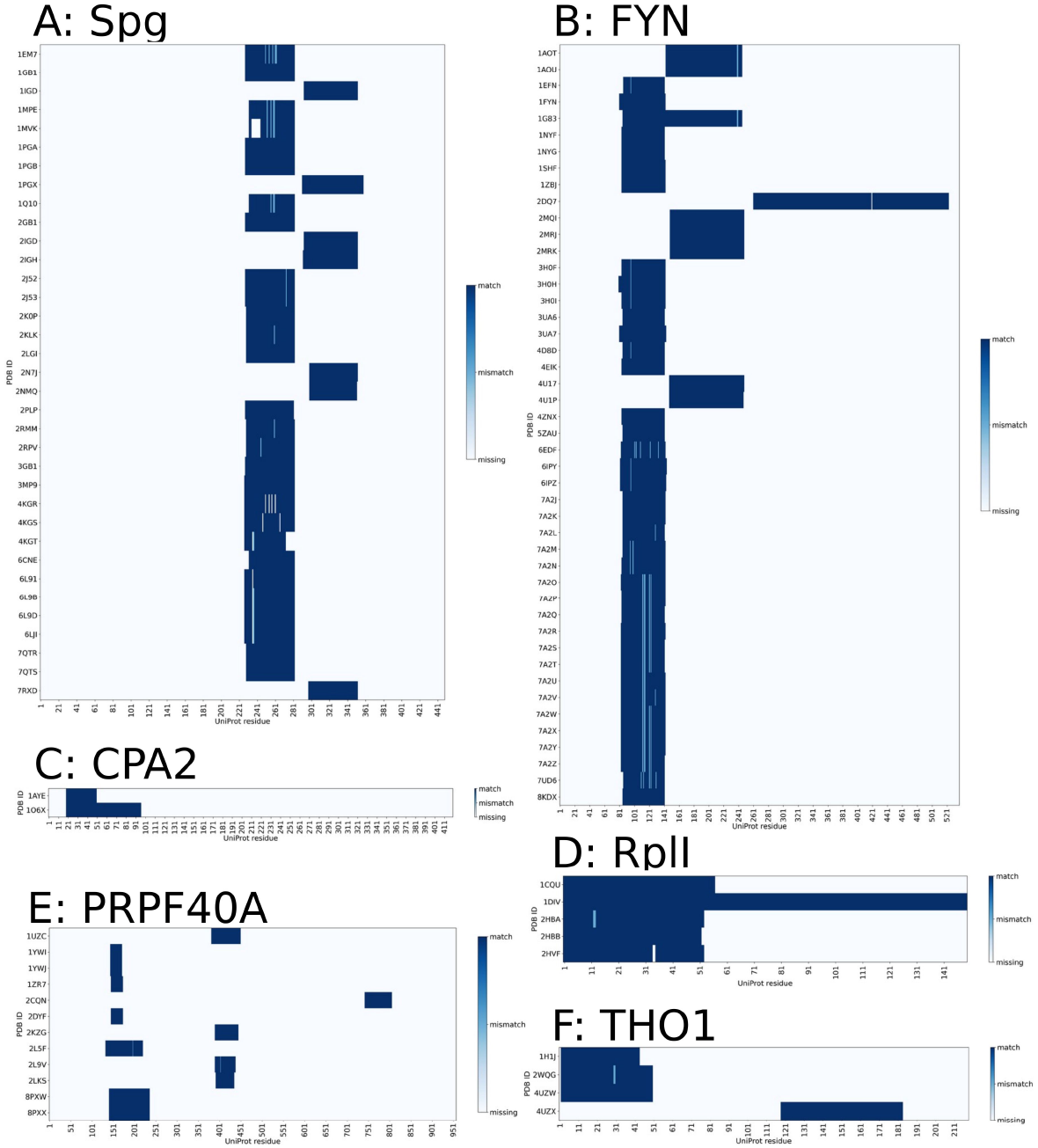
Alignment plots of other genes in this study

**Supplementary Figure 2.**
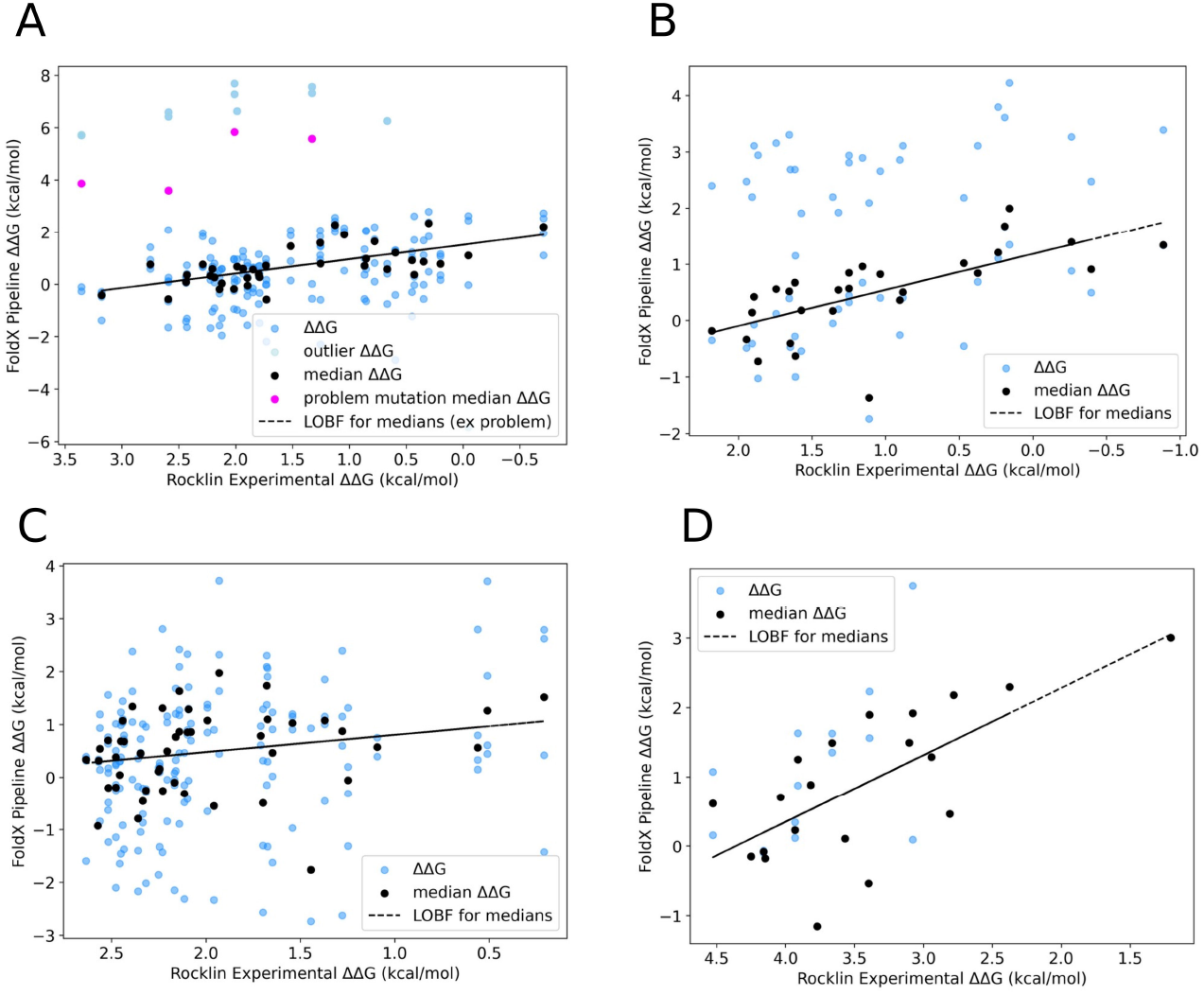
Regression analysis applied to other genes A) RPL1 B) THO1 C) PRPF40A D) CPA2

**Supplementary Figure 3.**
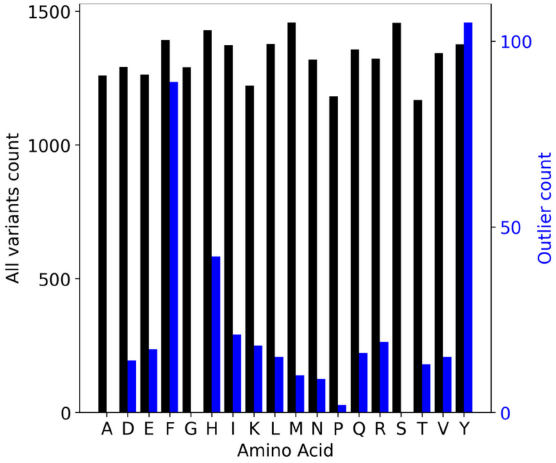
Frequency of outlier energy estimates aggregated by amino acid type.

**Supplementary Figure 4.**
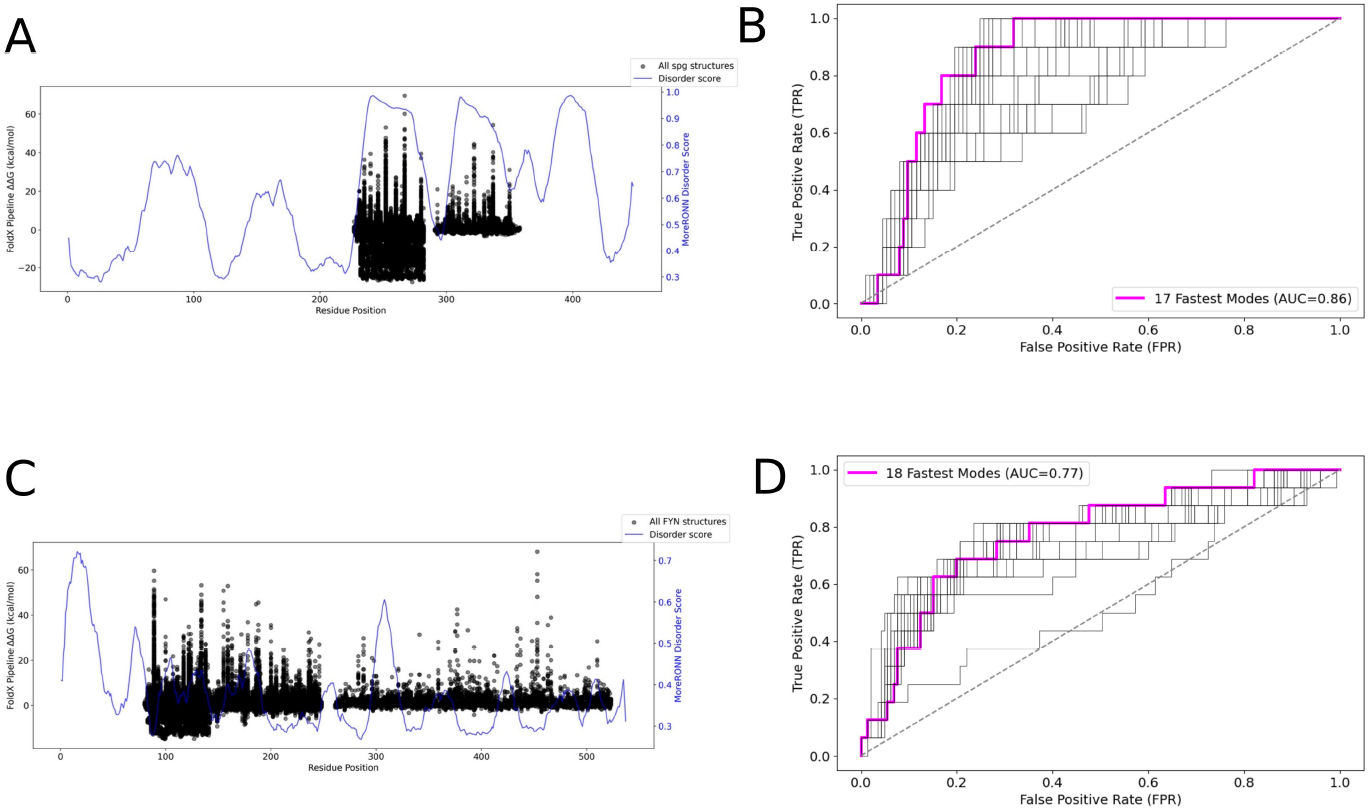
A, C) Distribution of FoldX pipeline ΔΔG energies across the full length of structures for SPG and FYN respectively with the MoreRONN disorder score overlaid. B, D) ROC plots for alternative numbers of top modes for SPG and FYN respectively. The multiple black curves represent the ROC curves for the first fastest mode up to the 30 fastest modes while the mode in magenta is the mode that maximises the area under the curve (AUC).

